# Rev-erbα heterozygosity produces a dose-dependent phenotypic advantage in mice

**DOI:** 10.1101/2019.12.30.890806

**Authors:** Ryan D. Welch, Cyrielle Billon, Thomas P. Burris, Colin A. Flaveny

## Abstract

Numerous mutational studies have demonstrated that circadian clock proteins regulate behavior and metabolism. *Nr1d1(Rev-erbα)* is a key regulator of circadian gene expression and a pleiotropic regulator of skeletal muscle homeostasis and lipid metabolism. Loss of *Rev-erbα* expression induces muscular atrophy, high adiposity, and metabolic syndrome in mice. Here we show that, unlike knockout mice, *Nr1d1* heterozygous mice are not susceptible to muscular atrophy and in fact paradoxically possess larger myofiber diameters and improved neuromuscular function, compared to wildtype mice. Heterozygous mice lacked dyslipidemia, a characteristic of *Nr1d1* knockout mice and displayed increased whole-body fatty-acid oxidation during periods of inactivity (light cycle). Heterozygous mice also exhibited higher rates of glucose uptake when fasted, and had elevated basal rates of gluconeogenesis compared to wildtype and knockout littermates. *Rev-erbα* ablation suppressed glycolysis and fatty acid-oxidation in white-adipose tissue (WAT), whereas partial *Rev-erbα* loss, curiously stimulated these processes. Our investigations revealed that *Rev-erbα* dose-dependently regulates glucose metabolism and fatty acid oxidation in WAT and muscle.

## Introduction

Circadian rhythmicity directs many aspects of behavior and metabolism [1]. The circadian clock is controlled by interconnected proteins that oscillate in activity and expression over a 24-hour cycle [2]. The central/master regulator of the internal clock is located within the Suprachiasmatic Nucleus (SCN) of the hypothalamus and integrates environmental light cues to synchronize peripheral oscillators in the liver, pancreas, and skeletal muscle (SkM) [1]. In addition to the central circadian clock, peripheral clocks can undergo autonomous oscillatory shifts in response to changes in nutrient availability without input from the central clock [3–6]. In all tissues the core molecular clock activators BMAL1 and CLOCK induce the transcription of circadian regulators PER and CRY that in turn inhibit CLOCK and BMAL1 activity, creating consistently robust diurnal oscillations [7]. This molecular clock has an auxiliary loop that is modulated by the nuclear receptors Retinoid-receptor like orphan receptor (ROR) and REV-ERBα/β (NR1D1/2) [8]. ROR, a transcriptional activator and REV-ERB a constitutive transcriptional repressor, reciprocally modulate each other’s activity by binding competitively to the same REV-ERB response elements (RevREs). Each receptor therefore oppositely regulate expression of the clock proteins BMAL1 and CLOCK, and therefore form an additional regulatory arm to the core molecular clock that enables more robust control of behavior in response to metabolic queues [8–10].

Disruptions of circadian rhythm is a well-characterized contributary factor to metabolic disorders in humans [1]. Mutational studies in mice have demonstrated that *Clock* deletion promotes obesity and metabolic syndrome characterized by hyperlipidemia, hyperleptinemia, hepatic steatosis, hyperglycemia, and deficient insulin production [11]. *Bmal1* mutant studies have demonstrated the importance of the molecular clocks in mitochondrial dynamics and type-2 diabetes protection [5, 12]. Additionally, the loss of *Bmal1* in mice is known to produce symptoms of advanced aging, including total body weight loss and sarcopenia [13]. REV-ERBα, like other circadian factors, is a key metabolic regulator of energy homeostasis. *Rev-erbα* deficient mice have higher basal glucose levels, enhanced fatty-acid synthesis-gene expression and display shifts in metabolic substrate preference in response to specific changes in light/dark cycles [14]. Moreover, *Rev-erbα* deficient mice display a reduction in lipid mobilization under fasted conditions and are consequently susceptible to high adiposity [14, 15].

REV-ERB has a pivotal role in muscle homeostasis as well. Previously, we have highlighted that REV-ERBα directs myogenesis through tethered interaction with the transcription complex Nuclear Factor-Y [16]. Moreover, inhibiting REV-ERB with a synthetic antagonist enhanced muscle repair in an acute injury model and slowed disease progression in a model of Duchenne’s muscular dystrophy [16, 17]. Loss of *Rev-Erbβ* in contrast to *Rev-Erbα* was also shown to stimulate skeletal muscle metabolism and fatty acid oxidation [18]. We have previously noted that *Nr1d1*^+/−^ mice had larger myofiber diameters compared to wild-type and *Nr1d1*^−/−^ mice, which implied that partial loss of REV-ERBα uniquely stimulated muscle hypertrophy and or inhibited muscle degradation. Heterozygous mice also showed an enhanced regenerative response to injury compared to wildtype mice [16]. *Rev-erbα* null mice are known to display elevated expression of mediators of muscle atrophy, impaired muscle function, and a progressive decline in myofiber size due to enhanced protein catabolism [19]. In order to decipher the metabolic underpinnings of this gene dose-effect we comprehensively characterized the metabolic phenotype of *Nr1d1*^+/−^ mice. We demonstrate that young *Nr1d1* heterozygous mice have larger myofibers and a greater lean mass than wildtype and knockout littermates. We also show that *Rev-erbα* heterozygosity enhanced lean mass composition and induced fatty oxidation in WAT adipose tissue as well as increased glucose sensitivity and enhances gluconeogenesis compared to *Rev-erbα* null and wildtype mice. Our results illustrate that there is a unique dose-threshold for REV-ERB regulation of metabolic activity. This discovery adds a new layer to the complexity of Rev-erb modulation of muscle homeostasis and lipid metabolism.

## Methods

### Mice

C57BL/6 (*Nr1d1*^+/+^) and B6.Cg-Nr1d1tm1Ven/LazJ (*Nr1d1*^+/−^ and *Nr1d1*^−/−^) mice were purchased from the Jackson laboratory. All mice were housed in a 12h-12h light-dark cycle, received a standard chow diet, and were allowed food and water *ad libitum*. At the conclusion to the study mice were sacrificed by CO_2_ asphyxiation followed by cervical dislocation. Mouse experimental procedures were approved by the Saint Louis University Institutional Animal Care and Use Committee (protocol#2474). Mouse sample size (n) was chosen using an α set at 0.05 a priori with a power of 80.

### NMR Body composition analysis

Nuclear magnetic resonance (NMR) analyses were conducted once a week on 6-week old *Nr1d1*^+/+^, *Nr1d1*^+/−^, and *Nr1d1*^−/−^ mice (n=7 per group) until 12 weeks of age by utilizing a Bruker BioSpin LF50 Body Composition Analyzer. Statistically significant differences in lean, and fat mass were determined by 2-way ANOVA with an alpha set at 0.05.

### H&E staining and cross-sectional area (CSA) analysis

Muscle samples (Tibialis anterior or gastrocnemius) were fixed overnight in 4% formalin and then embedded into paraffin. Paraffin embedded muscle sections where stained by a standard hematoxylin and eosin (H&E) protocol. Slide images were taken by a Leica MC120 HD attached to a Leica DM750 microscope. Cross-sectional area (CSA) of H&E stained myofibers where quantified by ImageJ. Six images from each experimental animal was utilized to calculate the average CSA per mouse. Images were taken at 10x and 40x for needed calculations.

### Grip strength

Hindlimb and forelimb grip strength were tested simultaneously with a digital force instrument (BIOSEB) as described previously [20]. Each (*Nr1d1*^+/+^, *Nr1d1*^+/−^ or *Nr1d1*^−/−^) mouse was subjected to 3 grip strength tests at 5 weeks and 11 weeks of age with a minimum of 10-minute rest intervals between each test. To reduce user-specific bias, two distinct researchers administered tests. Values from the two separate experimental groups were recorded and pooled. Statistical significance between the respective groups was determined using a student’s t-test with an alpha set at 0.05.

### Quantitative PCR

Total RNA was isolated from muscle tissue by using Trizol reagent (Invitrogen). Isolated RNA (1μg) was reverse-transcribed into cDNA using qScript cDNA Synthesis Kit (Quanta Biosciences, Inc.). Quantitative PCR (qPCR) was performed with SYBR Select Master Mix (Applied Biosystems) and cognate primers. Gene expression levels for mouse tissue and cells were normalized to *Gapdh*.

### Immunoblotting

Cells and muscle tissues were lysed by a RIPA with protease inhibitor buffer. The extracted proteins were flash frozen and stored at −80°C. The extracted proteins were separated by SDS-PAGE and transferred onto PVDF membranes. Immunoblot analyses were performed with standard procedures.

### Feeding study

Feeding behavior in 8-week-old *Nr1d1*^+/+^ and *Nr1d1*^+/−^ mice was assessed over 14 days using the BioDAQ episodic intake monitor. Mice were allowed food and water *ad libitum*. Data was collected and analyzed using BioDaq software. Statistically significant differences in daily food intake, daily water intake, and daily feeding events for each group of mice were determined using a student’s t-test with an alpha set as 0.05.

### Wheel-Running Activity

*Nr1d1*^+/+^ and *Nr1d1*^+/−^ (n = 6 per group) mice wheel-running activity were monitored for 4 weeks using wheel-running cages (Actimetrics) in circadian cabinets. For light/dark experiments, mice were kept on a strict 12h-12h light dark cycle mice. For dark/dark experiments, mice were kept in 24h darkness. The activity data was collected and actograms were generated using ClockLab (MatLab) software.

### Glucose tolerance test (GTT)

Mice were fasted for 6 hours prior to GTT. Each mouse was weighed and the needed dose of glucose calculated. Glucose was prepared in a glucose/PBS solution that contained 250mg/ml of glucose. Injection volume was calculated by BW(g) X 10uL of PBS. Fasting blood glucose was measured using a Glucometer (One Touch Ultra™) by tail bleeds. Blood glucose was measured at 15, 30, 60, and 120 minutes’ post glucose injections.

### Insulin tolerance test (ITT)

Mice were fasted for 6 hours prior to ITT. Each mouse was weighed and the needed dose of insulin calculated. Insulin was prepared in an insulin/PBS solution that contained 0.1U/ml of insulin. Injection volume was calculated by BW(g) X 10uL of PBS. Fasting blood glucose was measured using a Glucometer (One Touch Ultra™) by tail bleeds. Blood glucose was measured at 15, 30, 60, and 120 minutes’ post insulin injections.

### Pyruvate tolerance test (PTT)

Mice were fasted for 6 hours prior to PTT. Each mouse was weighed and the needed dose of pyruvate calculated. Pyruvate was prepared in a pyruvate/PBS solution that contained 100mg/ml of pyruvate. Injection volume was calculated by BW(g) X 10uL of PBS. Fasting blood glucose was measured using a Glucometer (One Touch Ultra™) by tail bleeds as above. Blood glucose was measured at 15, 30, 60, and 120 minutes’ post injections.

### Metabolic Cages

Whole body metabolic states were tested by indirect calorimetry in a Comprehensive Lab Animal Monitoring System (Columbus Instruments) for 3 days after 5 days of habituation at 22°C (room temperature) or 30°C (thermoneutrality). CLAMS operation and analysis were conducted as recommended by the manufacturer. Mice (12-16 weeks old) were single housed and light and feeding conditions mirrored home cage conditions. The respiratory exchange ratio (RER) was calculated by VCO2 and VO2 as determined by the light/dark cycle. VCO2, VO2, and heat production values were normalized to lean mass.

### Statistical analysis

Statistical significance was determined by subjecting mean values per group to students-t test unless otherwise specified. A value of p≤0.05 is considered statistically significant.

## Results

### Rev-erbα expression modulates body composition and muscle function

In accord with previous studies, we observed that heterozygous REV-ERBα mice in addition to possessing heightened regenerative capacity interestingly had a greater total average body mass than wildtype mice [16] (Fig 1A). Firstly, we quantified the expression of *Rev-erbα* in the SkM of *Nr1d1*^+/+^, *Nr1d1*^+/−^ and *Nr1d1*^−/−^ mice at CT8 and found that *Rev-erbα* gene expression was dose dependently linked to *Nr1d1* genotype as partial loss of *Rev-erbα* resulted in a reduction in REV-ERBα expression (Fig 1B). Importantly *Nr1d1*-heterozygous mice did not display a compensatory increase in *Nr1d2* expression. To further ascertain the role of REV-ERBα we assayed the body composition of *Rev-erbα* null (*Nr1d1*^−/−^), wild type (*Nr1d1*^+/+^) and *Rev-erbα* heterozygous (*Nr1d1*^+/−^) mice. Surprisingly, *Nr1d1*^+/−^ mice displayed a higher mean body mass compared to that of the wild type and knockout mice from 6 to 12 weeks of age (Fig 1C). Nuclear magnetic resonance (NMR)-based body composition analysis revealed that the enhanced size of heterozygous mice was primarily due to a stable elevation of total lean mass (Fig 1D). The percentage lean mass was greater in heterozygous mice up until 12 weeks of age where it then matched that of wildtype mice (Fig 1E). Over time knockout mice exhibited a stark decline in lean mass (Fig 1E). Knockout mice also showed increased adiposity with age, as shown previously [21], (Fig 1F-G). Conversely, *Nr1d1*^+/−^ mice adiposity was significantly lower than knockout and wildtype mice up until 12 weeks of age when it became indistinguishable from that of wildtype mice (Fig 1F-G). Interestingly, *Nr1d1*^+/−^ mice of 5 and 11-weeks of age also exhibited superior grip strength when compared to *Nr1d1*^+/+^ and *Nr1d1*^−/−^ mice (Fig 1H). These results hinted that Nr1d1 heterozygosity selectively increased lean mass levels and suppressed adipose tissue mass in young mice and suggested that partial *Nr1d1* loss may enhance neuromuscular function.

**Fig 1.**
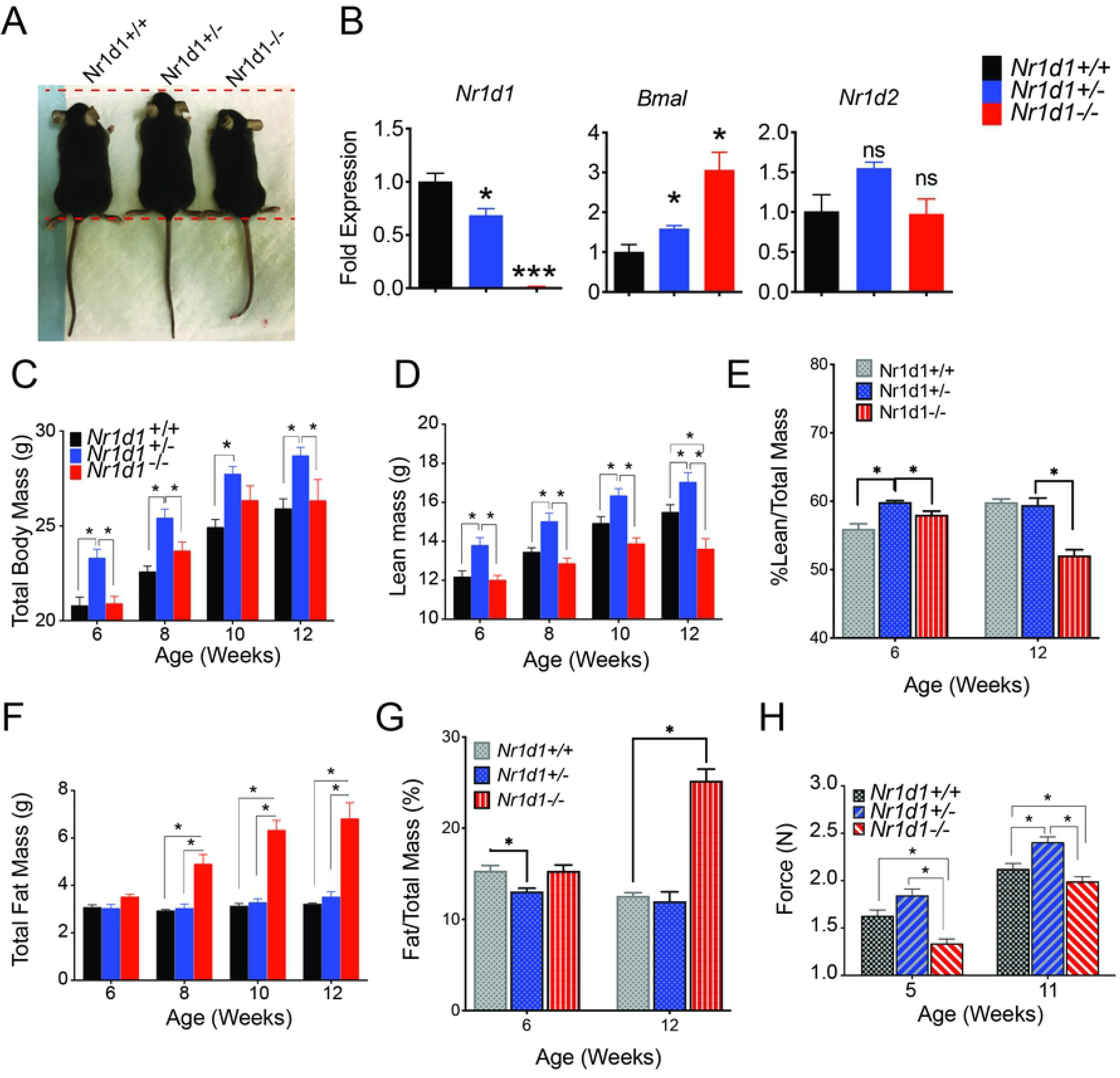
*Nr1d1* modulates body composition and grip strength in mice. (A) Representative sizes of *Nr1d1*^+/+^, *Nr1d1*^+/−^ and *Nr1d1*^−/−^ mice. Mice pictured are 6-week old males. (B) *Rev-erbα* and *Bmal1* mRNA transcripts in skeletal muscle of *Nr1d1*^+/+^, *Nr1d1*^+/−^ and *Nr1d1*^−/−^ at 5-7 weeks of age as determined by RT-qPCR (n = 4) (C) Total body weight of *Nr1d1*^+/+^, *Nr1d1*^+/−^ and *Nr1d1*^−/−^ at 6-12 weeks of age. (D) Total lean mass and (E) percent lean mass of *Nr1d1*^+/+^, *Nr1d1*^+/−^ and *Nr1d1*^−/−^ at 6-12 weeks of age (F) Fat mass and (G) percent fat mass of 6-12-week old *Nr1d1*^+/+^, *Nr1d1*^+/−^ and *Nr1d1*^−/−^ mice. Lean and fat mass were determined by using a Bruker BioSpin LF50 Body Composition Analyzer (n = 7). Statistically significant differences in lean, fat and fluid mass were determined by a 2-way ANOVA *p<0.05. (H). Forelimb and hindlimb grip strength of *Nr1d1*^+/+^, *Nr1d1*^+/−^ and *Nr1d1*^−/−^ mice. Grip strength was measured using a digital force instrument (BIOSEB) (n = 7). Grip strength data was analyzed using a student’s t-test *p<0.05.

### Nr1d1 heterozygous mice have normal circadian rhythms

It is known that full loss of *Rev-erbα* expression does not promote changes in wheel running activity in mice that are exposed to normal light/dark cycles [22, 23]. However, *Nr1d1*^−/−^ mice subjected to constant darkness display a shift in circadian activity, demonstrating that REV-ERBα is an important regulator of circadian behavior [22]. *Nr1d1*^+/−^ mice had normal circadian behavior when compared to wild-type mice in either normal light/dark cycles or in constant darkness, suggesting that the regulation of circadian function is preserved in the *Nr1d1*^+/−^ mice (Supplemental Fig 1A-B). Interestingly, this is despite heterozygous mice exhibitreduced expression of *Rev-erbα* and the significant increase in *Bmal1* expression (Fig 1B). These results collectively suggest that partial loss and homozygous deletion of Rev-erbα has contrasting effects on whole body composition that is not linked to changes in circadian regulation or feeding behavior.

### Heterozygous show enhanced myofiber size that is preserved after maturation

As previously mentioned, compared to wildtype littermates null mice show a rapid decline in lean mass after reaching maturity [16]. We analyzed the CSA of soleus muscle from *Nr1d1*^+/−^ and *Nr1d1*^−/−^ mice and found that, surprisingly, both knockout and heterozygous mice displayed significantly larger myofibers than wildtype littermates at 6-weeks-of age (Fig 2A). Interestingly, 15-week old *Nr1d1*^−/−^ mice showed progressive muscular atrophy while *Nr1d1*^+/−^ mice retained a larger myofiber size than that of wildtype mice (Fig 2B). To determine the underlying factors driving the distinct phenotypes we measured food/water intake and average number of feeding events in *Nr1d1*^+/−^ and *Nr1d1*^−/−^ mice. We observed no differences in the amount of food consumed, the frequency, or timing of feedings between *Nr1d1*^+/+^, *Nr1d1*^+/−^ and *Nr1d1*^−/−^ mice (Supplemental Fig 2A-C), suggesting *Nr1d1* heterozygosity did not influence lean mass through augmentation of feeding behavior. These results complement similar observations in *Nr1d1*^−/−^ mice that highlighted that complete loss of *Rev-erbα* did not produce changes in feeding activity and food intake when allowed food *ad libitum* [14, 15].

**Fig 2.**
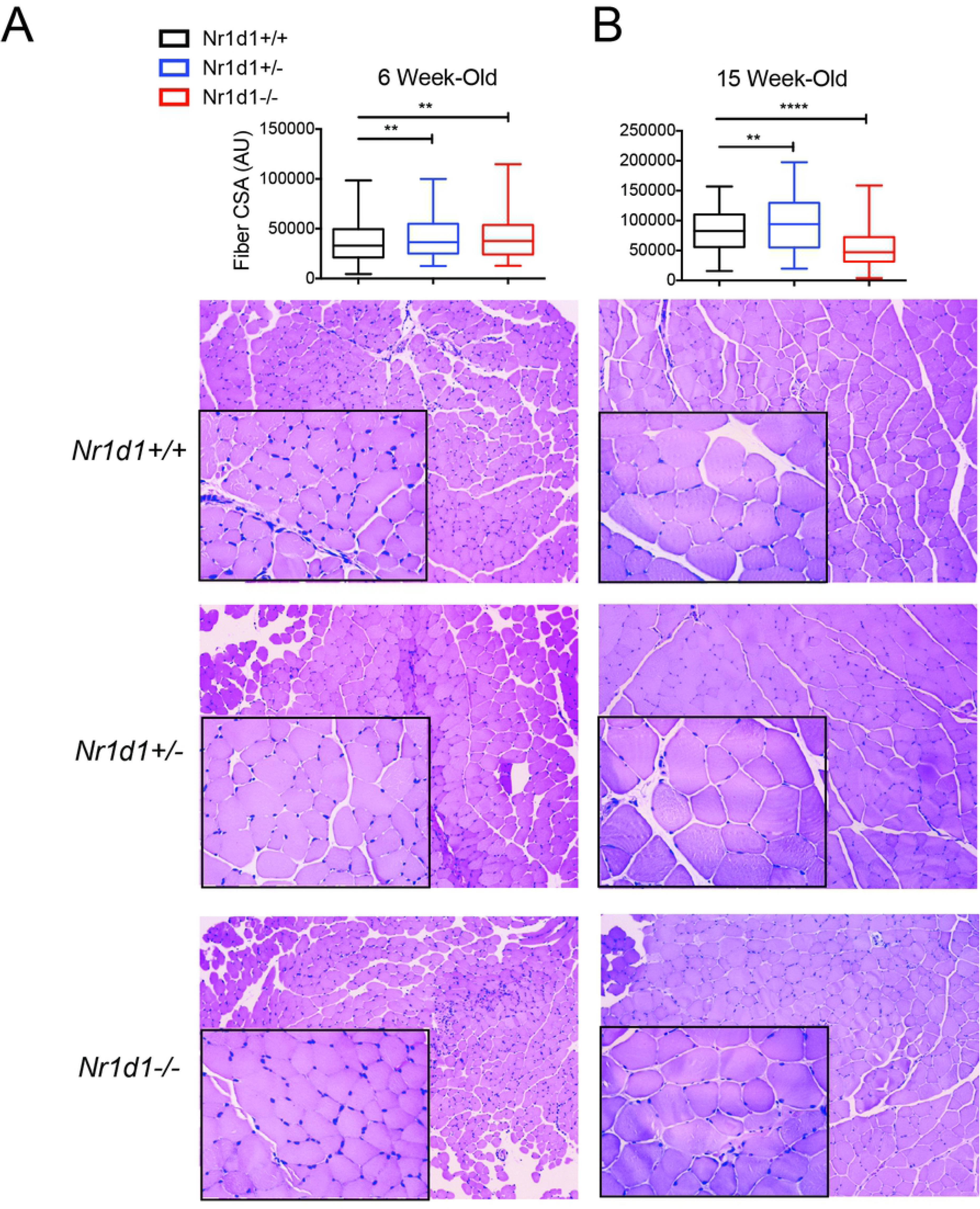
Reduced *Nr1d1* expression increases muscle fiber cross sectional area. (A-B) Quantification of myofiber cross-sectional area (CSA) (n = 4) and corresponding hematoxylin/eosin (H&E) stained tibia anterior muscle sections from A-5-7 and B-15 weeks old *Nr1d1*^+/+^, *Nr1d1*^+/−^ and *Nr1d1*^−/−^ mice. CSA was quantified using Image J software. **p<0.01 and ****p<0.0001 were determined by a student’s t-test. Data represented as box and whiskers plot.

### Partial loss of *Rev-erbα* expression does not alter catabolic or anabolic gene expression in skeletal muscle

To investigate why 15 weeks old *Nr1d1*^+/−^ and *Nr1d1*^−/−^ mice displayed varying lean mass levels we analyzed markers of cellular stress and protein metabolism. *Nr1d1*^+/−^ mice displayed no variation in gene expression of cellular stress markers *Ddit4, Map3k6, Map3k8, Map3k14, Tgif1, Mknk2, Junb*, and *Cebpb* (Supplemental Fig 3A). Ddit4 (or Redd1) is activated by DNA damage and inhibits mTOR signaling in SkM [24]. Interestingly the expression of the cell growth inhibitor *Ddit4* actually decreased in the *Nr1d1*^+/−^ mice. We also found no differences in catabolic and anabolic gene expression between the *Nr1d*^+/−^ and Nr1d1^+/+^ mice (Fig 3A-D). However, when *Nr1d1* expression was completely lost, cellular stress expression increased profoundly (Supplemental Fig 3B). We discovered by immunoblotting an increase in cleaved Caspase 3 levels in Rev-erbα-null mouse skeletal muscle, indicating enhanced protease activity (Supplemental Fig 3C). *Rev-erbα* null mice stimulated catabolic genes exclusively (Fig 3c) with no effect on anabolic expression (Fig 3D), in agreement with previous findings [19]. Further investigation revealed an increase in FOXO3A protein expression (Fig 3E) and STAT3 phosphorylation in the SkM of the *Nr1d1*^−/−^ mice (Fig 3F). In concordance with these findings, we detected a decrease in oxidative (slow) myosin heavy chains (Fig 3G) and glycolytic (fast) myosin heavy chains (Fig 3H) in the SkM of the *Nr1d1*^−/−^ mice. Overall, these data support the idea that *Rev-erbα* influences SkM protein stability through suppression of catabolic pathways particularly in older mice; a process that is disrupted in *Nr1d1*-null mice.

**Fig 3.**
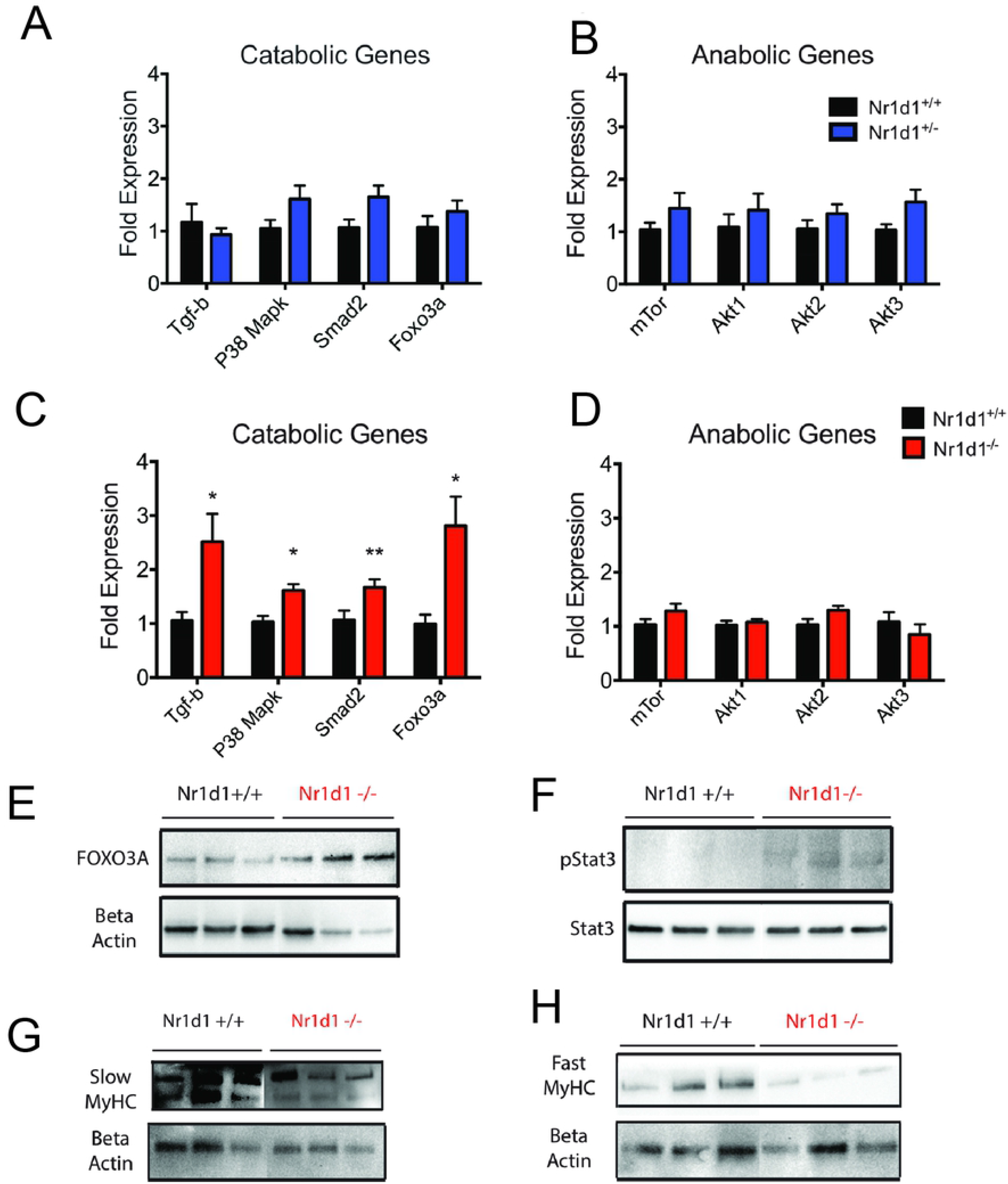
*Nr1d1* ablation induces catabolism of skeletal muscle partial Nr1d1 loss has no effect. (A-B) Catabolic and anabolic gene expression in 15 week old *Nr1d1*^+/−^ mice (n = 5-6). (C-D) Catabolic and anabolic gene expression in 15-week-old *Nr1d*^−/−^ mice (n = 5-6). Gene expression was determined by RT-qPCR. Immunoblot analysis of (E) FOXO3A, (F) p-STAT3/STAT3, (G) Slow and (H) Fast myosin heavy chains in the soleus muscle of 15 week old *Nr1d1*^−/−^ mice (n = 3). *p<0.05 and **p<0.01 were determined by a student’s t-test. RT-qPCR data are expressed as mean ± s.e.m.

### Decreased *Rev-erbα* expression induces fatty acid oxidation in mice

REV-ERBα is a regulator of metabolism and thermogenesis [25, 26]. However, whether *Nr1d1*^+/−^ mice exhibit a difference in whole body metabolism has not been previously elucidated. Adult *Nr1d1*^+/−^ mice, 12-14 weeks old, were placed in metabolic cages to quantify whole-body metabolism, using indirect calorimetry, and heat production. As expected, NMR-analysis revealed that *Nr1d1*^+/−^ mice had higher total body weight due to higher fat mass and increased lean mass relative to wildtype mice (Supplemental Fig 4A-C). Interestingly we observed no differences in VO2 and VCO2 either during the light or dark cycle (Fig 4A-B). Remarkably, we found a minor drop in the respiratory exchange ratio (RER) in heterozygous mice to a value of 0.88 during the light cycle, representing a modest increase in fatty acid substrate preference (Fig 4C). However, we found no difference in the substrate preference during the dark cycle in *Nr1d1*^+/−^ and *Nr1d1*^+/+^ mice (Fig 4D). Lastly, we found that *Nr1d1*^+/−^ mice displayed no changes in body heat production during the light cycle (Fig 4E). Still, during the dark cycle the Nr1d1-heterozygosity led to a mild increase in heat production (Fig 4F). These data suggest that even partially reducing *Rev-erbα* expression is sufficient to augment fatty acid metabolism and thermoregulation in a circadian dependent manner.

**Fig 4.**
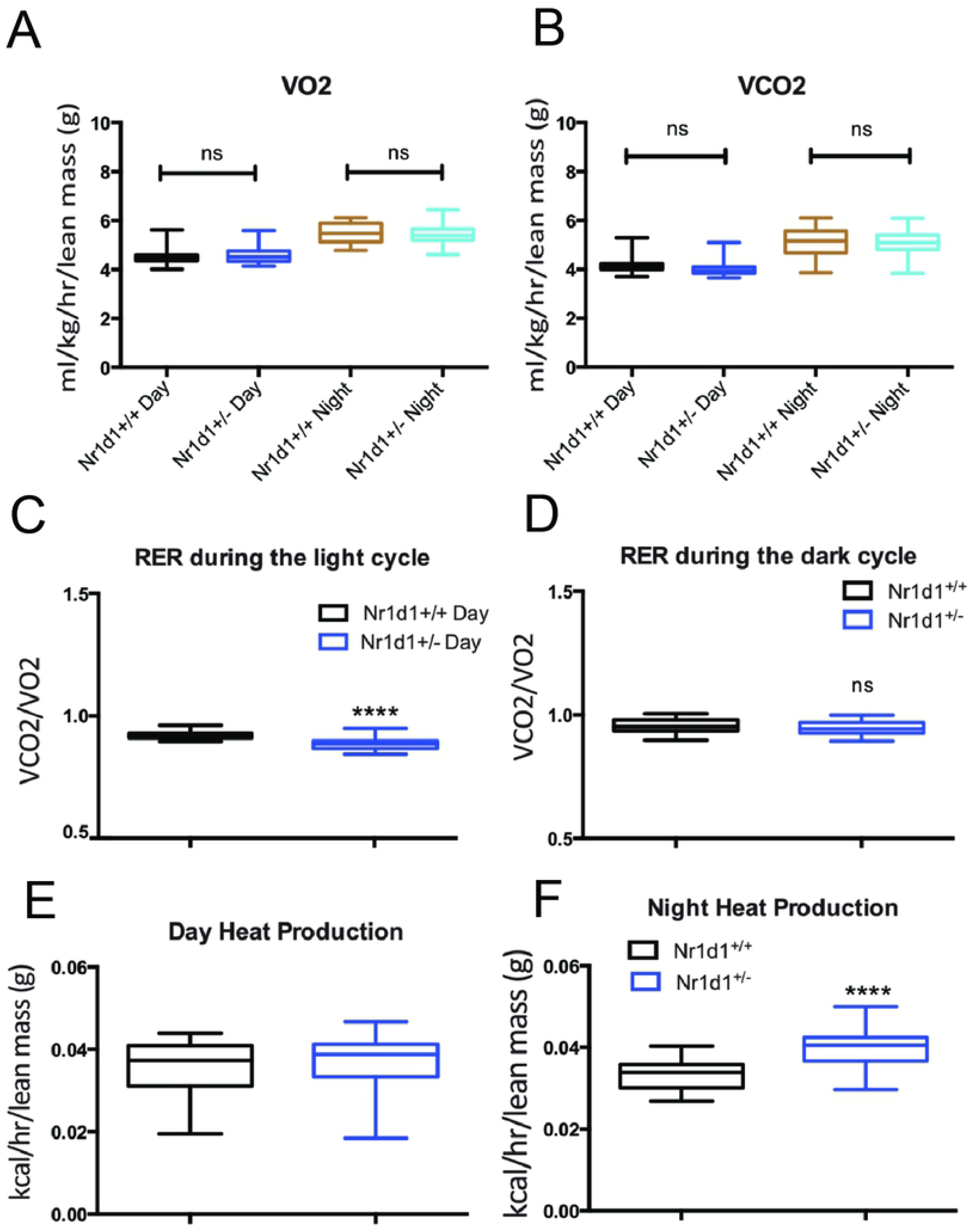
Reduced *Nr1d1* expression induces fatty acid utilization. All mice were kept on a 12:12 light/dark cycle at room temperature (n = 6). Whole body metabolism was determined by the use of Comprehensive Lab Animal Monitoring System (CLAMS). (A-B) Day or night analysis of VO2 and VCO2 of 12-14 weeks old *Nr1d1*^+/+^ *and Nr1d1*^+/−^ mice. (C-D) Respiratory Exchange Ratio (RER) of the *Nr1d1*^+/+^ and *Nr1d1*^+/−^ mice during the light and dark cycle. (E-F) Light and dark cycle heat production from the *Nr1d1*^+/+^ and *Nr1d1*^+/−^ mice. ****p<0.0001 was determined by a student’s t-test. VO2 and VCO2 were normalized to the lean mass of the animals. Data represented as box and whiskers plot.

### *Rev-erbα* heterozygosity expression increases glucose clearance independent of lean mass

To further characterize the metabolic effects of partial *Nr1d1* expression we decided to determine if wildtype and heterozygous mice displayed differences in glucose uptake, insulin sensitivity, and gluconeogenesis. To achieve this goal, we conducted a glucose tolerance test (GTT), an insulin tolerance test (ITT), and a pyruvate tolerance test (PTT) on *Nr1d1*^+/−^ and *Nr1d1*^+/+^ mice. We found no significant differences in the pre- and post-fast body weights of *Nr1d1*^+/−^ and *Nr1d1*^+/+^ mice (Supplemental Fig 5A). Importantly, heterozygous mice retained a higher percentage of lean mass and a similar percentage of fat mass to pre-fast wildtype mice (Supplemental Fig 5A-C). Interestingly, *Nr1d1* heterozygous mice had elevated blood glucose levels when fed compared to wildtype littermates (Fig 5A). However, fasted Nr1d1^+/−^ mice showed blood glucose levels equivalent to that of wildtype mice when fasted for 4 hours (Fig 5B). *Nr1d1*-heterozygous mice also displayed enhanced glucose clearance compared to wildtype mice (Fig 5C-D). However, heterozygous mice did not display any differences in insulin sensitivity compared to wildtype mice, as revealed by an insulin tolerance test (ITT) (Fig 5E). It should be noted that *Nr1d1*^−/−^ mice have not been shown to exhibit differences in glucose clearance and insulin sensitivity compared to wildtype mice, but do exhibit 10% higher resting blood glucose levels[14]. We also discovered that reduction of *Rev-erbα* expression increased gluconeogenesis as heterozygous mice significantly increased blood glucose levels in response to a bolus of pyruvate (Fig 5F). These results highlight the nuanced role of REV-ERBα in the regulation of glucose homeostasis and affirms the role of REV-ERBα as a mediator of gluconeogenesis in *vivo*.

**Fig 5.**
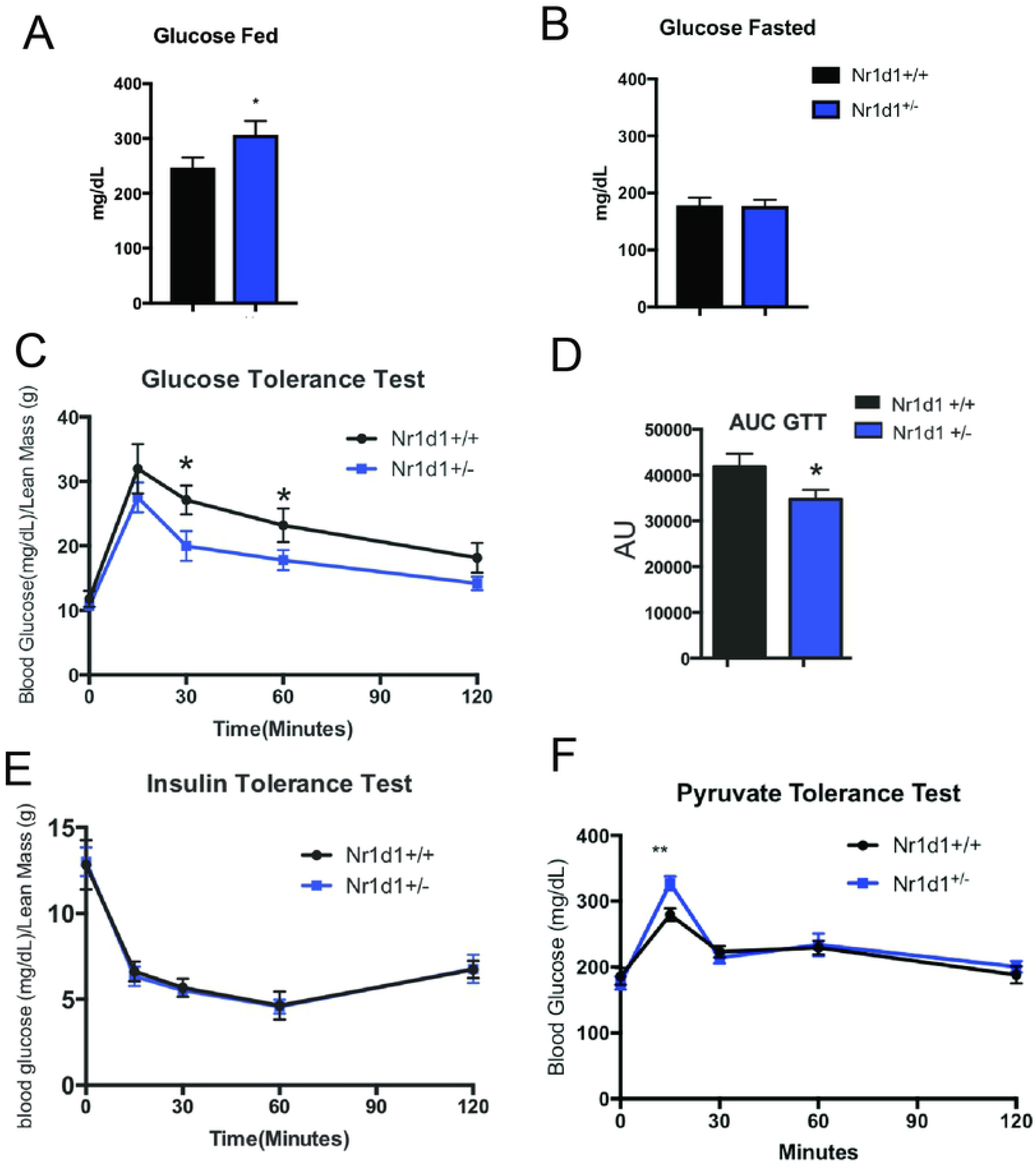
Partial loss of the *Nr1d1* gene promotes glucose clearance independent of lean mass levels. (A) Blood glucose levels in fed mice. (B) Blood glucose levels in 4 hours fasted mice. (C) Glucose tolerance test with (D) area under the curb analysis of *Nr1d1*^+/+^ *and Nr1d1*^+/−^ mice. (E) Insulin and (F) pyruvate tolerance test of *Nr1d1*^+/+^ and *Nr1d1*^+/−^ mice. Mice where 10-12 weeks of age when conducting GTT, 11-13 weeks of age when conducting the ITT, and 12-14 weeks of age when conducting the PTT (n =6). GTT and ITT were normalized to the lean mass of the animals. *p<0.05 and **p<0.01 were determined by student’s t-test. Data are expressed as mean ± s.e.m.

### Heterozygous *Rev-erbα* mice display a unique gene expression profile in distinct metabolic tissues

The partial reduction of *Rev-erbα* expression in *Nr1d1*^+/−^ mice induced a change in fatty acid oxidation and glucose metabolism that is not mirrored in *Rev-erbα* null mice [21, 27]. To gain a further understanding of the unique metabolic phenotype of *Nr1d1*^+/−^ mice, we probed whether partial *Rev-erbα* expression differentially modulated glycolytic and fatty acid oxidation in epididymal white adipose tissue (eWAT), brown adipose tissue (BAT), the liver, and soleus muscle. In the soleus muscle, we observed that *Rev-erbα* null mice showed an increase in fatty acid oxidation enzymes with a mild increase in expression of the glycolytic enzyme hexokinase 1 (*Hk1*) of the direct Rev-Erb target genes surveyed (Supplemental Fig 6A,C). Conversely, heterozygous mice displayed no differences in FAO gene expression (Supplemental Fig 6B), but exhibited an analogously mild decrease in the glycolytic genes *Hk1* and *Hk2* (Supplemental Fig 6D). These results highlighted that glycolytic and fatty-acid oxidation may not be differentially regulated in response to partial loss of Rev-Erbα.

In *Nr1d*^−/−^ mice livers RT-qPCR analysis revealed a drastic decrease in glycolytic enzyme expression with also a decrease in fatty acid oxidation genes (Fig 6A,B). Intriguingly, Nr1d1^+/−^ liver did not show changes in FAO enzyme expression but did also showed a more pronounced reduction in the expression of the glycolytic genes *Pfkfbp4, Hk1* and *Hk2* (Fig 6C,D), suggesting that *Nr1d1* heterozygosity may significantly influence hepatic glucose metabolism.

**Fig 6.**
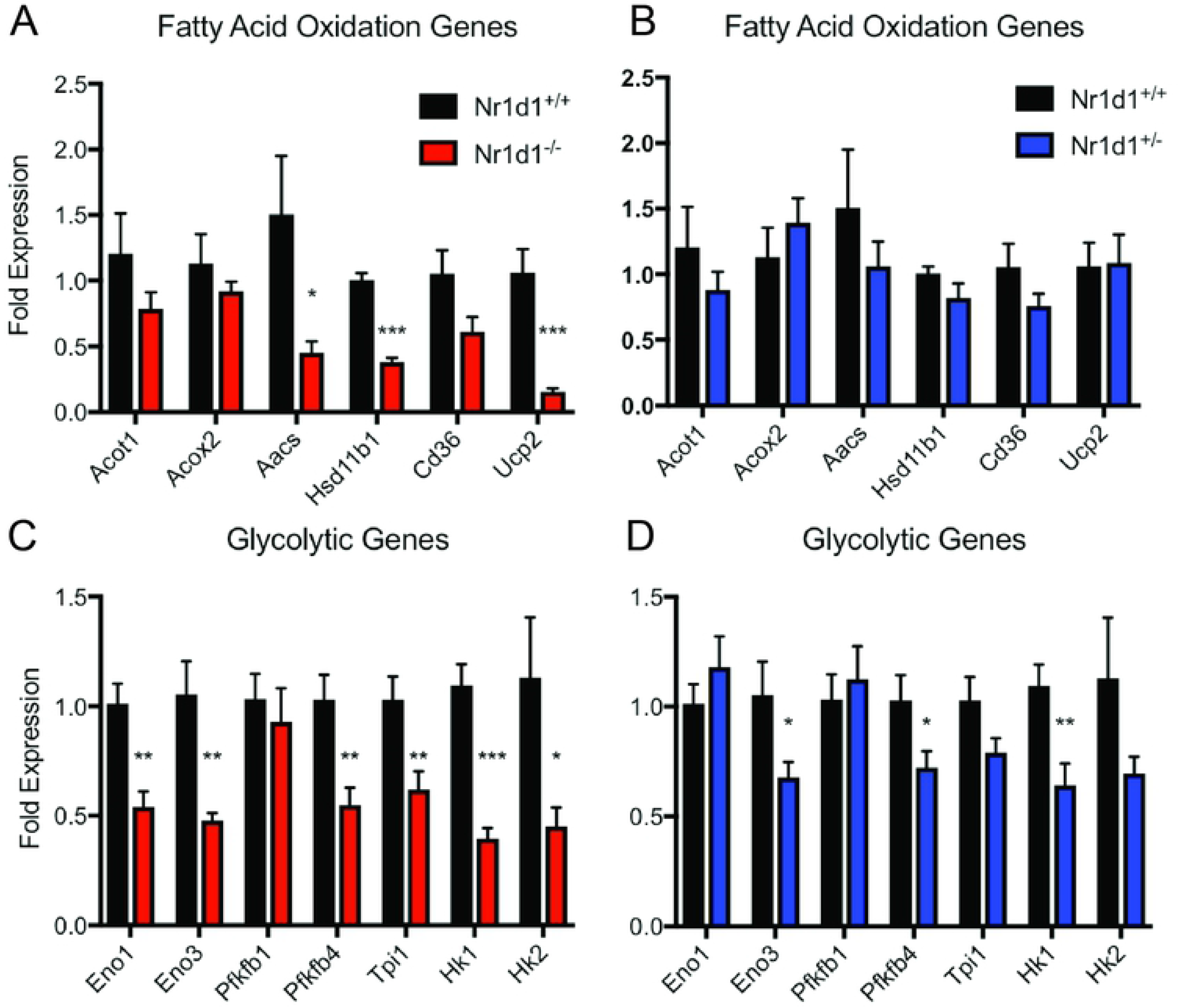
Heterozygous Nr1d1 liver display a reduction in glycolytic gene expression. Fold change of (A-B) fatty acid oxidation and (C-D) glycolytic gene expression in the livers of 15 weeks old *Nr1d1*^−/−^ and *Nr1d1*^+/−^ mice (n = 5-6). Gene expression was determined by RT-qPCR. *p<0.05, **p<0.01, and ***p<0.001 were determined by a student’s t-test. Data represented as a mean ± s.e.m. β

In BAT, Nr1d1^−/−^ mice uniquely exhibited a significant reduction in the glycolytic enzymes *Pfk1* and an induction of enolase 1 (*Eno1*) and triosphosphate-isomerase 1 (*Tpi1*) with partial *Rev-erbα* loss having no effect on these genes (Supplemental Fig 7A). Expression of *Hk1* and *Hk2*, however were similarly upregulated in both genotypes (Supplemental Fig 7A). Intriguingly only *Upc1* was upregulated in *Nr1d1*^−/−^ mice relative to wildtype and heterozygous mice, whereas other FAO factors such as *Ucp2*, *Ucp3* and *Cpt1a* being similarly downregulated in heterozygous and knockout mice versus wildtype mice (Supplemental Fig 7B). It is known that Rev-erbα null mice exhibit an elevated heat expenditure through increased *Ucp1* activity in BAT [28]. However, these results also highlighted that Rev-Erbα suppresses *Ucp2* and *Ucp3* gene expression in BAT.

In order to elucidate differences in Rev-Erbα mediated regulation of white adipose tissue metabolism, we also profiled glycolysis and FAO gene expression in WAT of *Nr1d1*^−/−^ and *Nr1d1*^+/−^ mice. We found that *Rev-erbα* produced an inverse expression pattern for FAO and glycolytic genes depending on the level of gene expression. In *Nr1d1* null mice eWAT displayed a decrease in glycolytic (*Pfk1*,*Pfkfbp4, Hk1 and Tpi1)* and FAO expression (*Ucp2*, *Aacs* and *Cpt1a*) yet these factors were elevated in *Nr1d1*^+/−^ mice (Fig 7A, B). Surprisingly, the fatty acid transporter *Cd36* was the only FAO factor downregulated in Nr1d1+/− mice compared to wildtype of knockouts (Fig 7A, B). These data suggest that decreased Rev-erbα expression stimulated, while in contrast complete loss of *Nr1d* paradoxically repressed eWAT metabolism. Suggesting the differences in metabolic phenotypes between the heterozygous and null mice may be due to a dose dependent differential regulation of glycolytic and fatty acid gene expression in WAT tissue.

**Fig 7.**
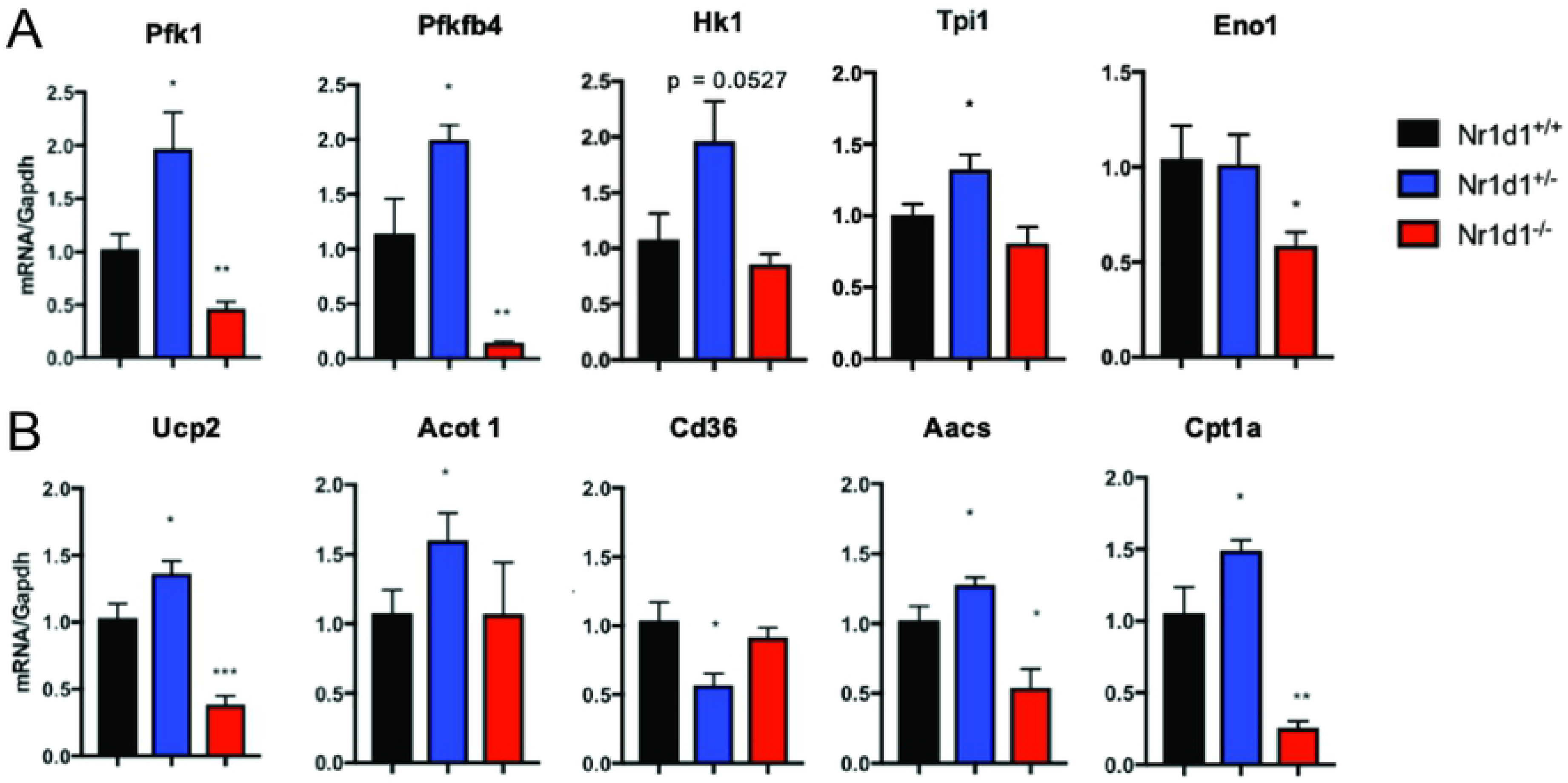
Heterozygous *Nr1d1* epididymal white adipose tissue (eWAT) exhibit enhanced metabolic gene expression. Fold change of (A) glycolytic and (B) fatty acid oxidation gene expression in the eWAT of 15 weeks old *Nr1d1*^−/−^ and *Nr1d1*^+/−^ mice (n = 5 - 6). Gene expression was determined by RT-qPCR. *p<0.05 and **p<0.01 were determined by two tailed-student’s t-test. Data represented as a mean ± s.e.m.

## Discussion

Circadian proteins are key regulators of metabolism and dictate susceptibility to metabolic syndromes [6, 11, 14, 27]. The nuclear receptor REV-ERBα is a key repressor of the molecular clock and modulates metabolic and developmental pathways. Our study wished to thoroughly understand how *Rev-erbα* expression levels impacts metabolism and muscle morphology due to a previous observation that *Nr1d1* heterozygous mice have superior muscle fiber size [16]. In this study we illustrate that *Nr1d1*-heterozygosity increased glucose clearance, fatty acid utilization, gluconeogenesis, and also boosted lean mass accumulation without generating higher adiposity in chow-fed mice.

Distinct differences in phenotype has been rarely reported with a beneficial effect of heterozygosity when compared to null animals. For example, this “Goldilocks Effect” has been described in *Ctrf* knock out mice resulting in decreased compliance and increased resistance in the lung and effectively mirroring the cystic fibrosis [29]. However, when mice were heterozygous for the mutated *Ctrf* allele it did not induce a classical gene dosage effect that was predicted by the phenotype. Surprisingly, *Ctrf*^+/−^ mice displayed decreased resistance and enhanced compliancy within the lung [29]. Resulting in a heterozygous advantage over both the null and wild type animals. This study, however, is the first to highlight the underlying contributory factors that drive the discordance between *Nr1d1* heterozygous and knockout mice. Our investigations aver that complete loss of REV-ERBα has deleterious effects on muscle homeostasis through enhanced metabolism of fatty acids in muscle and depressed glucose utilization and FAO in eWAT. Conversely, heterozygous mice displayed a bias toward fatty acid oxidation in eWAT during inactive periods, possibly driven by enhanced glycolytic and FAO enzyme expression. Heterozygous mice also interestingly exhibited enhanced thermogenesis during times of activity and enhanced glucose and pyruvate tolerance. Most importantly we show that the negative effects of REV-ERBα loss in muscle may in part be driven by disregulated muscle catabolism via FOXO3A and Stat3 activation in an age-dependent manner.

The mechanism(s) through which *Nr1d1* heterozygosity boosts lean mass levels at an early age is not clear. Our data demonstrates that *Nr1d1* null mice display significant gains in fat mass with a concomitant increase in muscular atrophy and an associated loss in muscle function. Intriguingly, heterozygous *Nr1d1* mice exhibit no changes in adiposity, showed increased grip strength, and enhanced muscle mass without significant changes in the expression of anabolic or catabolic factors in the skeletal muscle. Myogenic programs are highly active during neonatal growth and are one of the primary drivers in the development of the musculature [30]. We have found that myogenic gene expression is enhanced in *Nr1d1* heterozygous mice only when subjected to skeletal muscle injury, suggesting the *Nr1d1* heterozygosity is beneficial in skeletal muscle only under a promyogenic environment [16]. In mature post mitotic skeletal muscle we found no differences in metabolic gene expression among heterozygous versus wiltype mice, implying that reduced Rev-erbα expression may not impact mature skeletal muscle energetics. Therefore, it is plausible that mature *Nr1d1*^+/−^ mice experience the benefits of increased lean mass and strength due to key events early on in muscle development that facilitates and exponentially amplified increase in muscle mass by full maturity. *Rev-erbβ* is also highly expressed in post mitotic skeletal muscle and in some cases can act as an axillary component to REV-ERBα-mediated gene repression [23, 31]. Both *Rev-erbα* and *Rev-erbβ* expression is drastically reduced during myoblast differentiation [32, 33]. This study suggests that there may be an ideal threshold at which REV-ERBα expression facilitates elevated fatty oxidation in eWAT and lean mass accumulation. Amador and colleagues have suggested that REV-ERBβ in contrast to REV-ERBα knockouts may also exhibit muscle hypertrophy through enhanced food intake [18]. Here we show that partial loss of Rev-erbα expression alone sufficiently enhances lean mass without influence feeding behavior or activity. Importantly, differences between the *Rev-erbα* heterozygote and wildtype phenotype have only been assessed in skeletal muscle undergoing developmental growth or regenerative repair. Whether partial loss recapitulates the deleterious effects of Rev-erbα ablation on endurance and mitochondrial function should be the subject of future investigations.

Collectively our results indicate that Nr1d1 heterozygous mice exhibit a drastic departure from the phenotype of knockout mice. The data indicates that partial loss of *Nr1d1* gene does not produce deleterious effects and effectively augments metabolic substrate preference. This aligns well with our previous studies in which we showed that REV-ERBα inhibition is an effective approach for pharmacologically stimulating muscle repair [16, 17].

Polymorphisms in the *NR1D1* gene has been associated with obesity in various human populations [34, 35]. To our knowledge a correlation between lean mass and REV-ERBα SNPs has not been reported. New studies investigations that venture beyond assessments of fat mass levels and designed to probe the effect of *NR1D1* SNPs on whole-body metabolism may further shed light on the role of *REV-ERBα* expression in metabolic regulation in humans.

